# Structural basis of Norrin recognition by Lgr4/5/6

**DOI:** 10.1101/2022.09.29.510199

**Authors:** Tao-Hsin Chang, Karl Harlos, Fu-Lien Hsieh, E. Yvonne Jones

## Abstract

Lgr4/5/6 have been identified as stem cell markers and canonical Wnt signalling enhancers via complexing with R-spondin and Rnf43/Znrf3. Lgr4/5/6 have been reported to interact with Norrin, a cystine-knot growth factor that activates Wnt signalling via binding to Frizzled4 cysteine-rich domain (Fz4_CRD_). Norrin–Fz4 signalling axis regulates central nervous system vascularization. However, the molecular basis of Norrin and Lgr4/5/6 recognition remains lacking. Here, we present the structure of Norrin in complex with an ectodomain variant of Lgr4 (Norrin–Lgr4_ECD_), revealing a 2:2 stoichiometry. Our analyses show that residues of Lgr4/5/6_ECD_ for Norrin binding are highly conserved. Structural comparisons of Norrin–Lgr4_ECD_ with Rspo1– Lgr4_ECD_ and Norrin–Fz4_CRD_ complexes reveal that Norrin binding site on Lgr4 overlaps with Rspo1 binding site on Lgr4 and Fz4 binding site on Norrin overlaps with Lgr4 binding site on Norrin. The present work opens new avenues to elucidate the function of Norrin–Lgr4/5/6 signalling axis.

## Introduction

Canonical Wnt/β-catenin signalling – activation by secreted lipid-modified Wnt with Frizzled (Fz) receptor and LRP5/6 co-receptor – plays a central role in embryonic development and tissue homeostasis^1,2^. Abnormal Wnt signalling results in oncogenesis and degeneration^3,4^. To finely balance signalling activity, Wnt signalling is therefore tightly regulated by diverse cell surface and secreted proteins^5–7^.

One of those Wnt signalling regulations is to ubiquitylate Fz receptors for degradation by Rnf43 and Znrf3 (two transmembrane E3 ubiquitin ligases) that result in the attenuation of Wnt signalling^8,9^ (Fig. 7a). Furthermore, the ubiquitylation function of Rnf43/Znrf3 can be neutralized by complexing with the leucine-rich repeat-containing G protein-coupled receptor homolog 4,5, and 6 (Lgr4/5/6) and the secreted R-spondin (four homologs, Rspo1-4) proteins for membrane clearance via the endocytic process (Fig. 7b), thereby resulting in more Fz receptors on the cell surface and stronger Wnt signalling^10–13^. At the level of molecular interaction (Fig. 7b), Rspo – consisting of two cysteine-rich repeats (Fu1 and Fu2) and a C-terminal thrombospondin-related domain (TSP) and a positively charged tail – uses the Fu1 domain to interact with the extracellular domains of Rnf43/Znrf3 and Fu2 domain to interact with the extracellular domain (ECD) of Lgr4/5/6^14–19^.

Lgr4/5/6 are evolutionarily conserved and related to GPCRs for thyroid-stimulating hormone, follicle-stimulating hormone and luteinizing hormone^20,21^. In light of function, the currently accepted view of Lgr4/5/6 is to finetune Wnt signal strength for stem cell function via complexing with Rspo and Rnf43/Znrf3^10–13^. However, various lines of evidence suggest that Lgr4/5/6 can also activate the cAMP-dependent pathway through Gαs/Gαq signalling and the transforming growth factor-β signalling^22–25^.

Intriguingly, Lgr4/5/6 was reported to interact with a cystine-knot growth factor, Norrin which was encoded by *Norrie Disease Protein* (*NDP*) gene^26^. Mutations in the *NDP* gene have been characterized in Norrie disease (ND) and Familial exudative vitreoretinopathy (FEVR) – both of which result in hypovascularization of the retina, retinal detachment, and vision loss^27,28^. In function, Norrin has been well known to be the ligand binding to Fz4 cysteine-rich domain (Fz4_CRD_) that activates canonical Wnt/β-catenin signalling together with LRP5/6 co-receptor and Tetraspanin-12 co-activator^29,30^. Norrin mediated Wnt/β-catenin signalling has been reported to control central nervous system (CNS) angiogenesis and blood-brain-barrier (BBB) integrity^31^. However, the molecular basis of Norrin and Lgr4 recognition is unknown. Here, we revealed the complex structure of Norrin–Lgr4_ECD_ that adopts a 2:2 stoichiometry. Structure-guided mutagenesis in combination with protein-protein interaction assay defines the basis of Norrin and Lgr4/5/6 recognition. Our structural comparisons – Lgr4_ECD_ vs. Fz4_CRD_ for Norrin recognition and Norrin vs. Rspo for Lgr4/5/6 recognition – provide critical insights into divergent competition and cooperation of ligand and receptor recognitions for cellular signalling regulation. These insights would inform the investigation of the Norrin and Lgr4/5/6 signalling axis in angiogenesis and neurogenesis and the design of novel therapeutic interventions for modulating Norrin and Wnt signalling pathways.

## Results

### Lgr4/5/6 recognize Norrin via the extracellular domain

To determine whether Norrin interacts with the ECD of Lgr4/5/6, we probed the protein-protein interactions using the colourimetric alkaline phosphatase (AP) reaction-based method (Fig. 1a), using a reported probe of AP-Norrin (AP was fused to the N-terminus of Norrin)^29,32^. The Fc fusion baits of Lgr4/5/6_ECD_ with the C-terminal Avi-tag were expressed and biotinylated *in vivo* in human embryonic kidney (HEK293) cells and captured on a streptavidin-coated micro-well. As shown in Fig. 1b, Norrin binds to Lgr4/5/6_ECD_, in agreement with a previous work showing that Norrin bound to transfected cells with full-length Lgr4/5/6 on the cell surface^26^.

**Figure 1.**
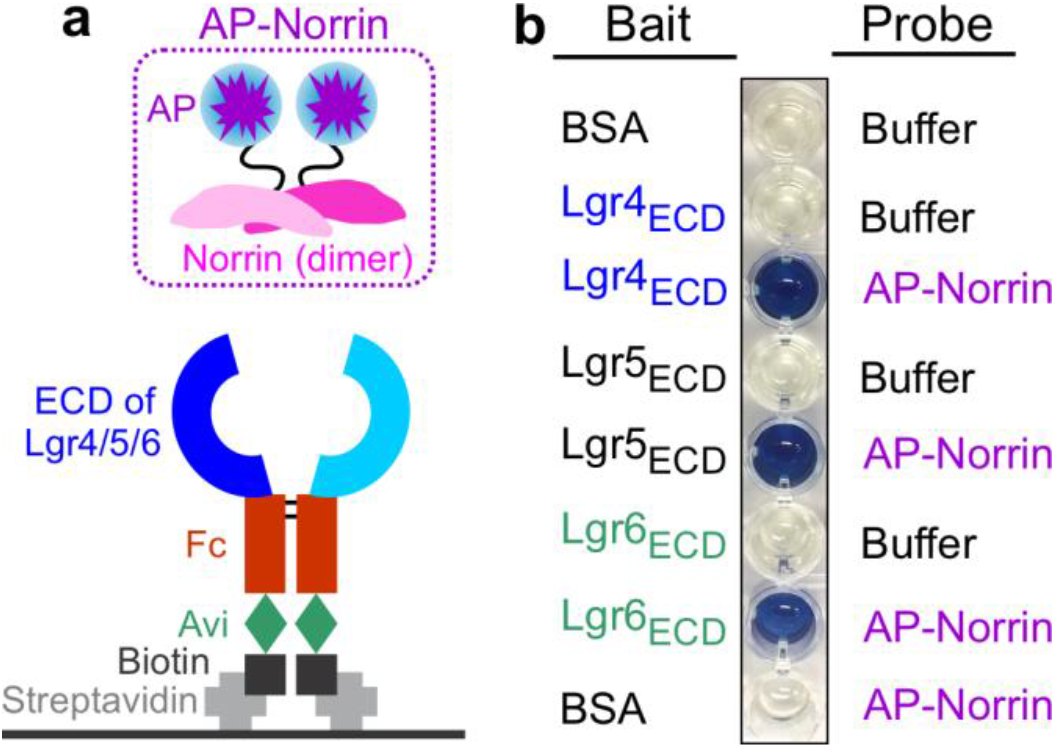
Norrin binds to the extracellular domain (ECD) of Lgr4/5/6. **(a)** Schematic diagram of AP-based protein-protein interaction assay. The biotinylated Avi-tag at the C-terminus of Fc fusion baits (Lgr4_ECD_, Lgr5_ECD_, and Lgr6_ECD_) were immobilized on streptavidin-coated wells and incubated with AP-Norrin (probe). **(b)** Bound AP-Norrin was visualized with BluePhos phosphatase substrate solution (a colourimetric AP reaction). BSA is used as a negative control.

### Production of Norrin–Lgr4_ECD_ complex proteins

To dissect the molecular mechanism of how the Lgr4/5/6_ECD_ recognizes Norrin, we selected Lgr4_ECD_ to complex with Norrin for structural and functional characterizations. Because of the challenge of obtaining stable Norrin–Lgr4_ECD_ complex proteins, we used the following strategies. Firstly, human Norrin was fused with an N-terminal small ubiquitin-like modified (SUMO) to boost Norrin expression in mammalian cells (Supplementary Fig. S1a), as we reported previously^33^. Secondly, to prevent protein aggregation of the ECD of Lgr4 ^18,34^, we adopted the hybrid LRR method^34^: the N-terminal LRR (residues 28-396) of Lgr4 fused with the C-terminus region (residues 131-200) of the hagfish variable lymphocyte receptor (VLR) protein (Supplementary Fig. S1a). We refer this variant of chimaera Lgr4_ECD_ to hereafter as Lgr4_ECD_. Thirdly, we co-expressed the SUMO fusion Norrin together with the Lgr4_ECD_ with a C-terminal mVenus fusion protein in HEK293 cells (Supplementary Fig. S1a). The secreted complex was purified and removed the fusion proteins (SUMO and mVenus). Norrin forms a stable protein complex with Lgr4_ECD_ during purification (Supplementary Fig. S1b). To investigate the stoichiometry of complex formation in solution, we used size-exclusion chromatography coupled with multi-angle light scattering (SEC-MALS) to demonstrate the formation of a 2:2 complex (Supplementary Fig. S1c and Table S1).

### The overall structure of Norrin in complex with Lgr4_ECD_

The obtained Norrin–Lgr4_ECD_ complex was subject to crystallization trials using *in situ* deglycosylation method^35^, allowing a complete data to be collected and processed to 3.05 Å resolution with anisotropic correction (Supplementary Table 2). Next, we determined the complex structure of Norrin–Lgr4_ECD_ by the molecular replacement method using Norrin and Lgr4_ECD_ as the search models (Fig. 2 and Supplementary Fig. S2). As expected from the SEC-MALS result (Supplementary Fig. S1c), the structure of the Norrin–Lgr4_ECD_ complex revealed a 2:2 stoichiometry, that two Lgr4_ECD_ interact one-to-one with a separate Norrin chain without contacting each other (Fig. 2). Norrin adopts a very elongated dimer (stabilized by three intermolecular disulphide bonds, Cys93-Cys95, Cys95-Cys93, and Cys131-Cys131) in a head-to-tail fashion (Fig. 2 and Supplementary Fig. S3), presenting distinct features from other cystine-knot growth factors e.g. transforming growth factor and bone morphogenetic protein ^33,36^. Lgr4_ECD_ adopts a horseshoe-like architecture with a twisted solenoid structure (Fig. 2), which is composed of the central LRR (1-15) flanked by the LRR N-terminal domain (LRRNT) and the C-terminal VLR.

**Figure 2.**
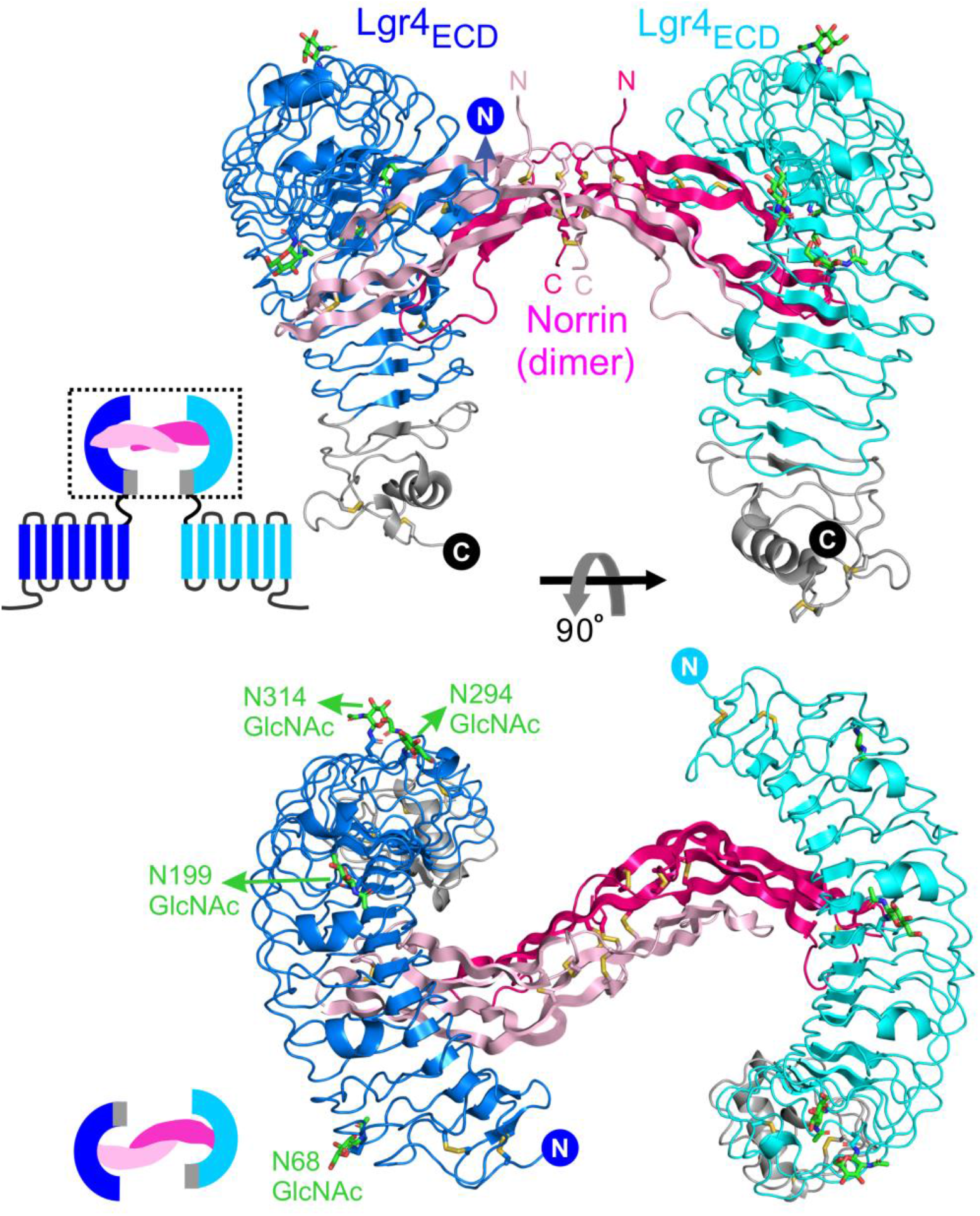
The structural basis of Norrin–Lgr4_ECD_ complex. The structure of Norrin (pink and magenta) in a 2:2 complex with Lgr4_ECD_ (blue and cyan) in two views. N-linked glycosylation sites and associated glycans are shown as green sticks. The insets show cartoon models of Norrin and Lgr4 signalling complex in different orientations. The hagfish VLR fusion proteins are coloured in grey. The N- and C-termini are labelled.

### Structural and functional characterization of the binding interface

In the binding interface, Norrin uses the regions (β1 to β2 and β5 to β6) far from the centre dimerization region to contact the concave surface of Lgr4_ECD_, which includes the LRRNT and LRR (1-10), through electrostatic and hydrophobic interactions (Fig. 3 and Supplementary Table 3). The binding interface between Norrin and Lgr4_ECD_ buries a surface area of ~825Å^2^. The residues of Lgr4 for Norrin binding are highly conserved in Lgr5 and Lgr6 (Supplementary Fig. S4), in agreement with that all Lgr4/5/6 can recognize Norrin (Fig. 1). Notably, several Norrin residues for Lgr4 binding are reported as disease-associated mutation sites, identified from genetic retinal disorders e.g., ND and FEVR (Fig. 3 and Supplementary Table 4). To further examine the binding interface, we mutated the residues of Lgr4 for Norrin binding by introducing N-linked glycosylation sites (Fig. 4a). Lgr4 mutations (W159N/D161T, H207N/H209T, and D231N/N233T) abolish the interaction of Lgr4 with Norrin (Fig. 4b).

**Figure 3.**
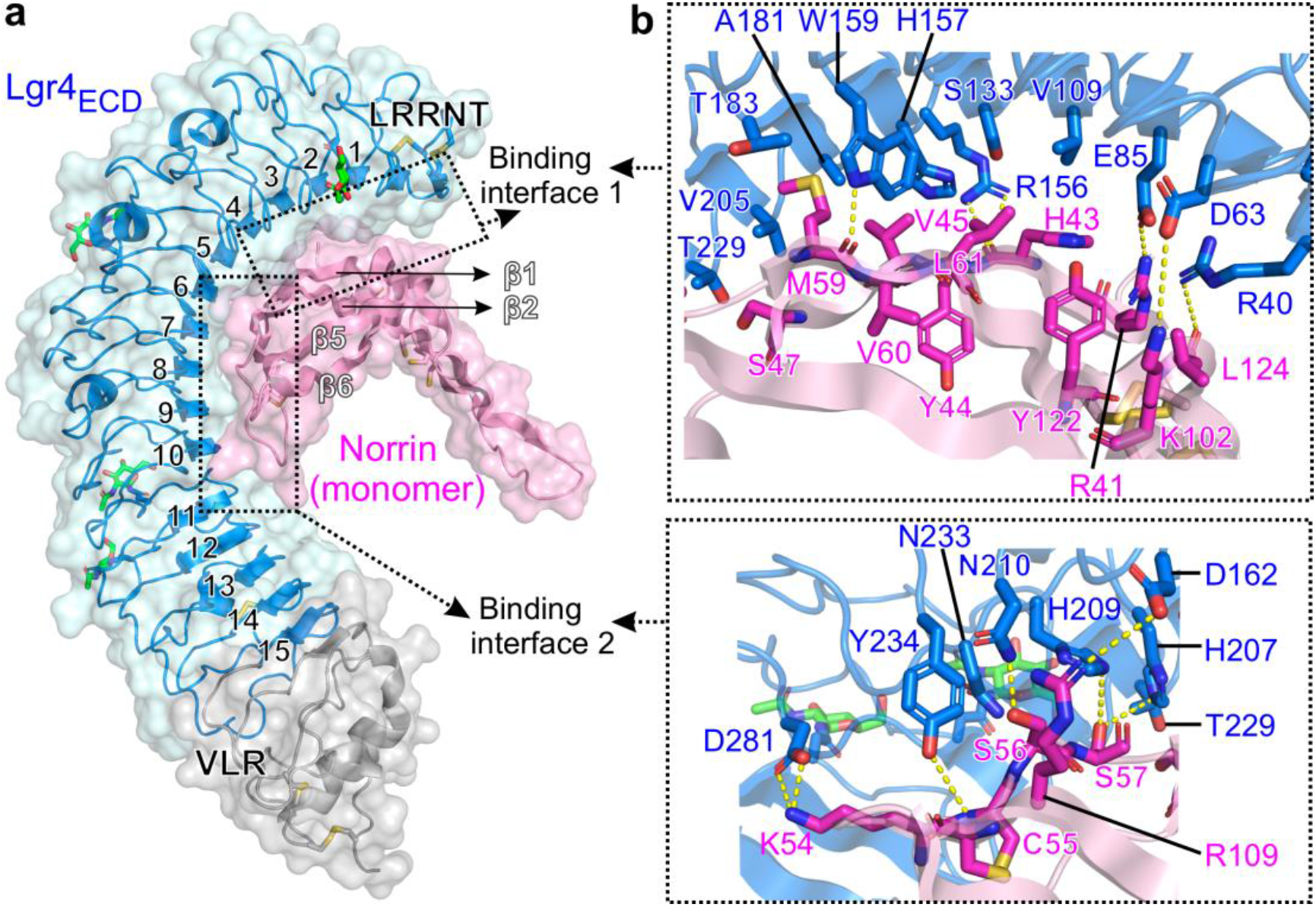
Structural details of binding interfaces in Norrin–Lgr4_ECD_ complex. **(a)** Side-view of complex shown in semi-transparent surface and ribbon representation of Norrin and Lgr4_ECD_. The central 15 LRR repeats of Lgr4 are labelled. The two binding interfaces are marked with dotted boxes. **(b)** A close-up view of the binding interfaces. Residues involved in binding are shown as sticks and labelled (Norrin in magenta and Lgr4_ECD_ in blue). Dotted lines denote hydrogen bonds. N-linked glycans are shown as green sticks.

**Figure 4.**
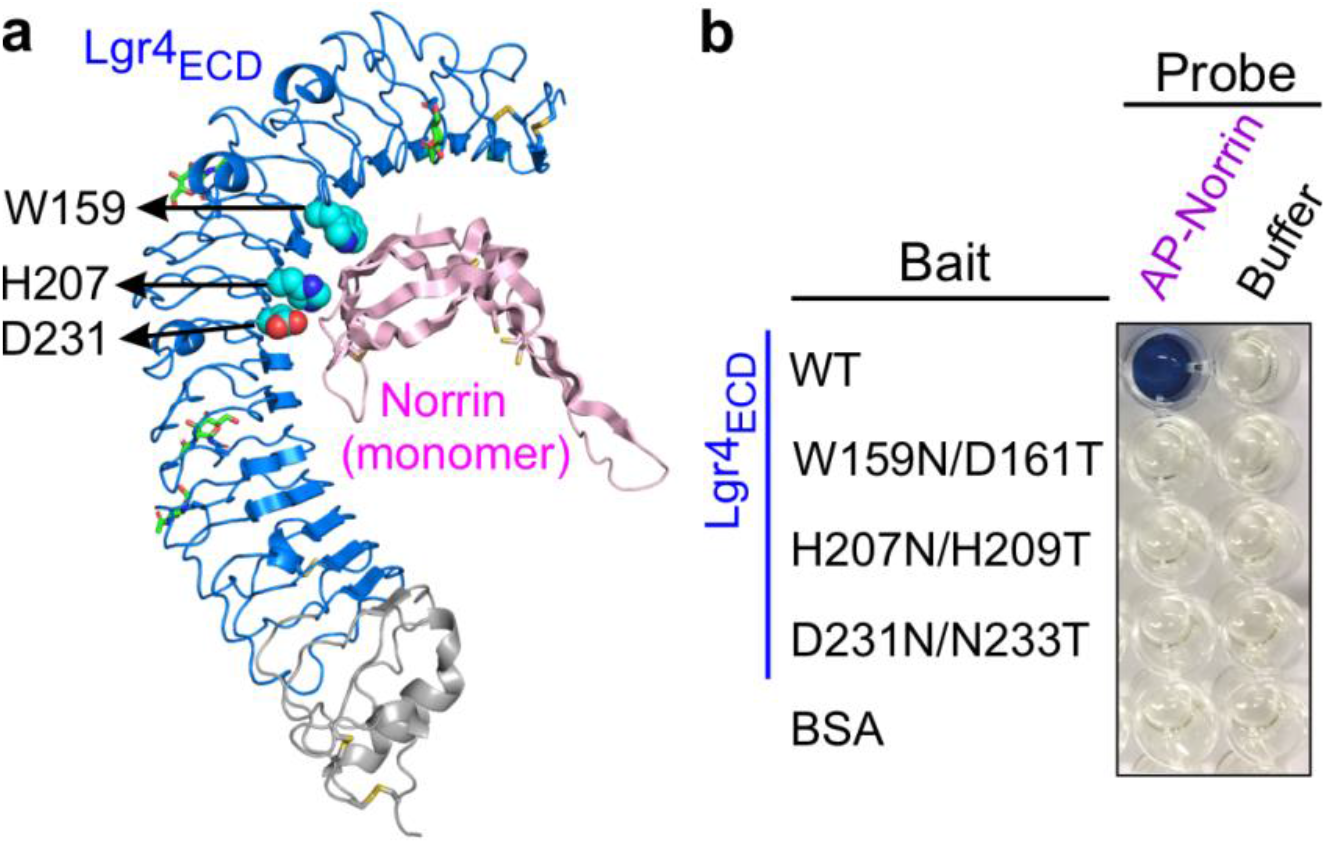
Functional characterization of Norrin and Lgr4/5/6 interactions. **(a)** Residues of Lgr4 in the binding interface for mutagenesis are shown as spheres (atom colouring: cyan, carbon; blue, nitrogen; red, oxygen). N-linked glycans are shown as sticks. **(b)** AP-based protein-protein interaction assay of Lgr4_ECD_ mutants and Norrin interactions.

### Structural comparison of Norrin–Lgr4 with Rspo1–Lgr4 and Norrin–Fz4

Lgr4/5/6 are well characterized for the recognition of Rspo proteins^14–16,18,37^. To compare the binding modes of Lgr4/5/6 for Rspo and Norrin binding, we superimposed Rspo1–Lgr4_ECD_ with Norrin–Lgr4_ECD_ (Fig. 5a). Intriguingly, the position of Rspo1 for Lgr4 binding generally overlaps with Norrin (Fig. 5a), although Rspo1 and Norrin present distinct protein folds. Furthermore, our detailed binding interface analyses also revealed that the Norrin binding region on the Lgr4 surface largely overlaps with the Rspo1 binding region (Fig. 6a). In other words, Norrin and Rspo can compete with each other for Lgr4/5/6 binding. Additionally, the LRRNT of Lgr4 folds down toward the centre concaveness upon Norrin binding (Fig. 5a), in agreement with that the LRRNT of Lgr4 contribute to Norrin binding (Fig. 3a).

**Figure 5:**
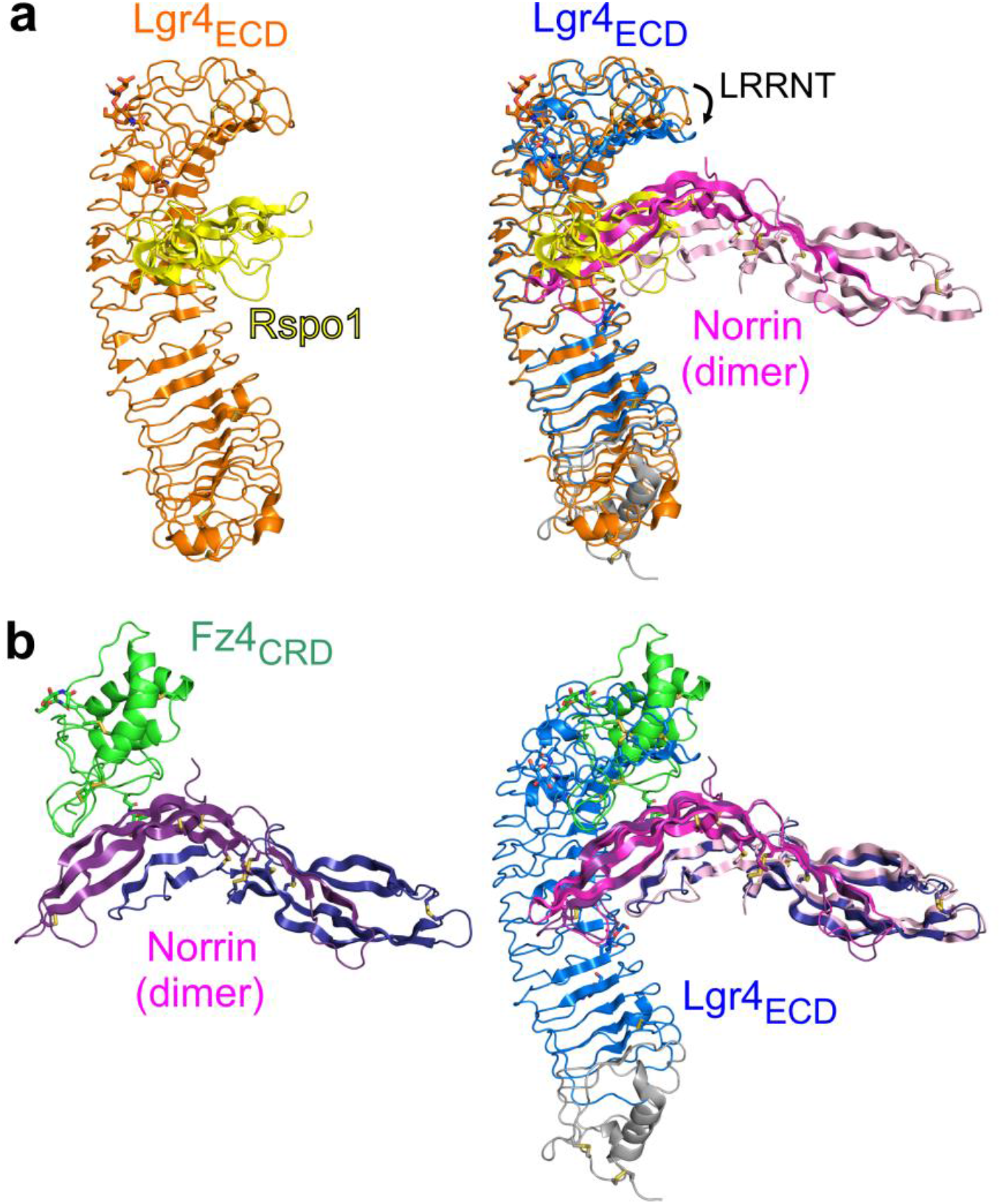
Structural comparison of Norrin–Lgr4_ECD_ with Rspo1–Lgr4_ECD_ and Norrin–Fz4_CRD_. **(a)** The complex structure of Rspo1–Lgr4_ECD_ (PDB code 4KT1) and superimposition of Rspo1–Lgr4_ECD_ with Norrin–Lgr4_ECD_ based on Lgr4_ECD_. **(b)** The complex structure of Norrin–Fz4_CRD_ (PDB code 5BQC) and superimposition of Norrin–Fz4_CRD_ with Norrin–Lgr4_ECD_ based on Norrin as a reference.

**Figure 6.**
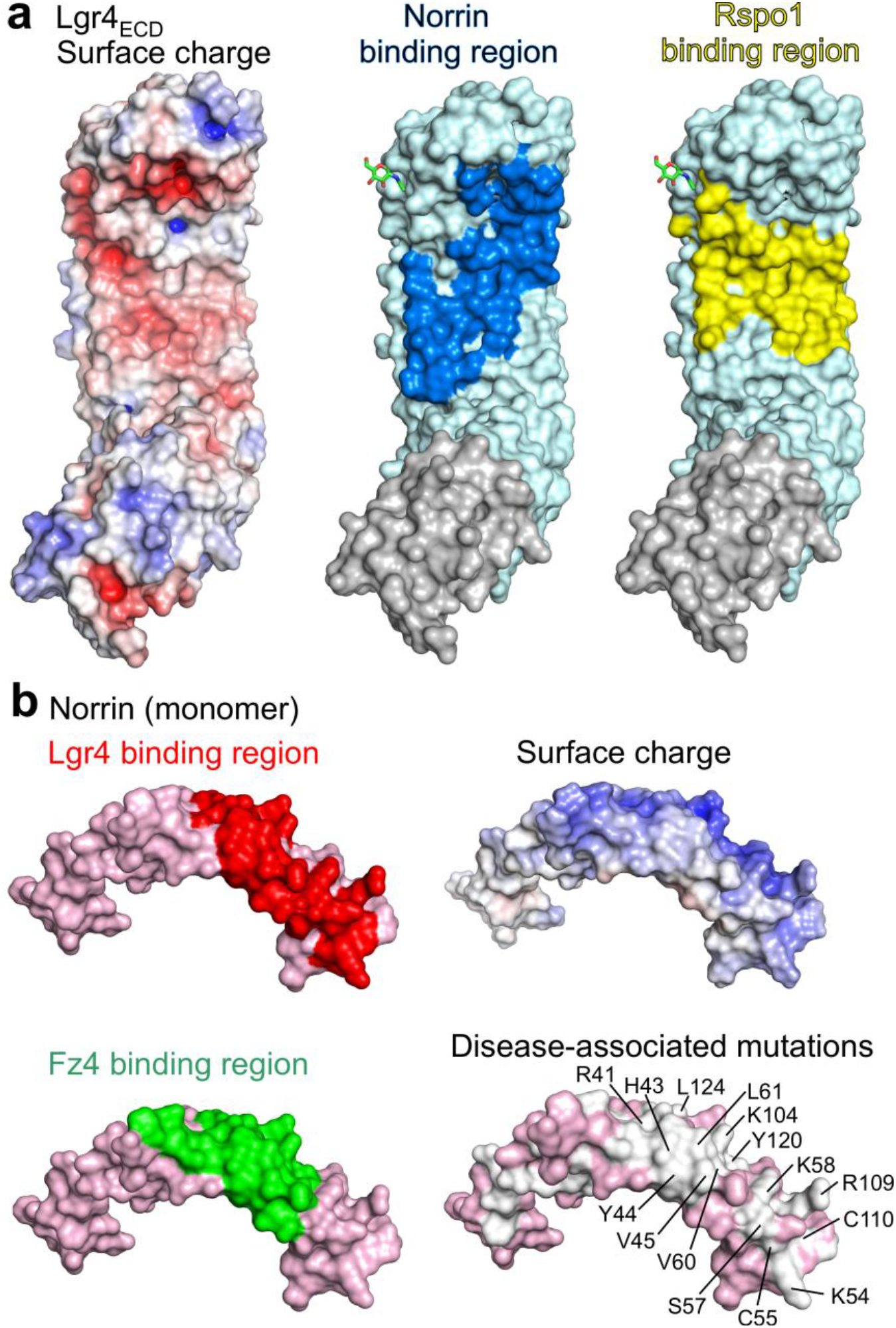
Structural analyses of Norrin and Rspo1 binding regions on Lgr4 and Fz4 and Lgr4 binding regions on Norrin. **(a)** Electrostatic surface potential of Lgr4_ECD_ is coloured from red (acidic; −7 *k*_b_*T*/_*e*_c__) to blue (basic; 7 *k*_b_*T*/_*e*_c__). Surface representations of Lgr4_ECD_ are coloured cyan and grey for VLR fusion protein. Interface residues of Lgr4 for Norrin binding and Rspo1 binding are coloured blue and yellow, respectively. **(b)** Surface representations of Norrin (pink). Interface residues of Norrin for Lgr4 binding and Fz4 binding are coloured red and green, respectively. Electrostatic surface potential of Lgr4_ECD_ is coloured from red (acidic; −7 *k*_b_*T*/_*e*_c__) to blue (basic; 7 *k*_b_*T*/_*e*_c__). Disease-associated mutations mapped onto the surface of Norrin are coloured grey and mutation sites overlapped with interface residues for Lgr4 binding are labelled.

Intriguingly, Norrin has been reported to activate the canonical Wnt signalling through binding to the Fz4 cysteine-rich domain (Fz4_CRD_)^29,33,38^. To dissect the molecular mechanism of Norrin for Fz4 and Lgr4 binding, we superimposed Norrin–Fz4_CRD_ with Norrin–Lgr4_ECD_ (Fig. 5b). Notably, Fz4_CRD_ clashes sterically with Lgr4_ECD_ upon Norrin binding (Fig. 5b), consistent with the Lgr4 binding region on the Norrin surface partially overlapping with Fz4 binding region (Fig. 6b). Norrin adopts conserved conformations upon Fz4 and Lgr4 binding, suggesting that Norrin utilize a rigid structural fold to interact with Fz4 and Lgr4. Interestingly, Norrin residues for Lgr4 and Fz4 binding overlap with disease-associated mutations (Fig. 6b and Supplementary Table 4).

## Discussion

The present study provides several advances for our understanding of Norrin/Wnt signalling and ligand–receptor recognition.

Firstly, Norrin binds not only to Fz4_CRD_ but also to the ECD of Lgr4/5/6 (Fig. 1), in agreement with our structure and sequence analyses revealing that residues on Lgr4/5/6 for Norrin binding are highly conserved (Supplementary Fig. S4). As Norrin–Fz4 signalling axis is well characterized in CNS angiogenesis and BBB integrity as well as vision deficit in ND and FEVR, it is particularly interesting to ask what is the physiological function of Norrin–Lgr4/5/6 signaling axis (Fig 7c)? Intriguingly, Norrin has been found to protect damaged retinal neurons ^39^ and induce retinal progenitor proliferation^40^. Recently, Miller *et al*., further showed that Norrin and Lgr6 are enriched in astrocyte and glia, and both modulate neuronal activity, dendritic growth, and spine formation^41^. The treatment of Norrin can improve neuronal connectivity and strength^41^. Interestingly, the spine and dendrite deficits have been found in epilepsy and Alzheimer’s disease^42,43^. Furthermore, Orre et al., reported that transcriptomic Norrin is downregulated in astrocytes of the Alzheimer’s disease model^44^. Indeed, human brain RNA-seq data^45^ also revealed that Norrin and Lgr4 have markedly high expression in astrocytes, compared to neuron and endothelial cells (Supplementary Fig. S5). Collectively, do these observations explain clinical characterizations of cognitive and behavioural deficits in ND and FEVR? It will need to dissect whether the abnormal Norrin–Lgr4/5/6 signalling axis results in this pathological progress.

**Figure 7.**
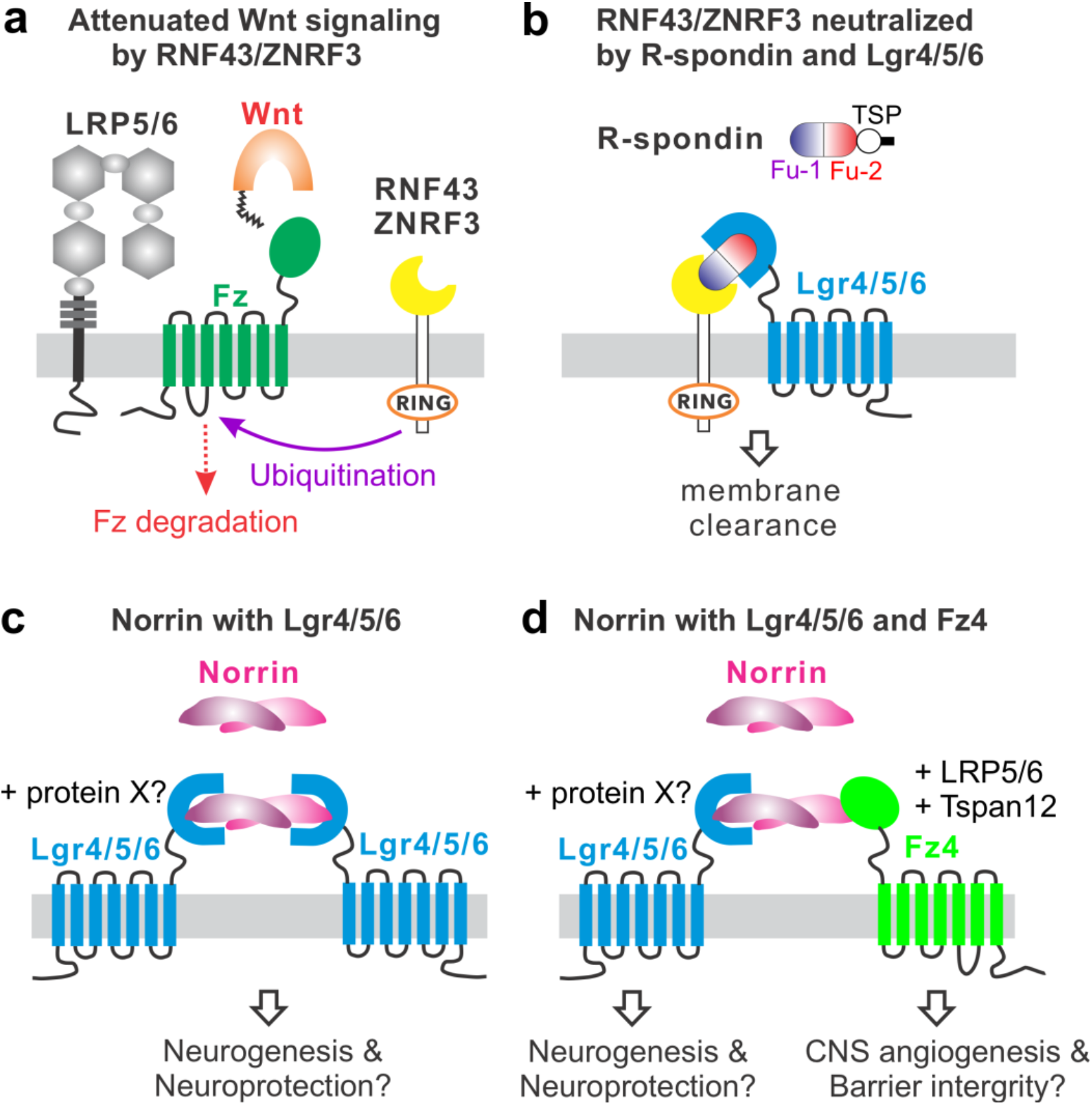
Regulation of Wnt signalling and proposed signalling modules. **(a)** Wnt signalling can be attenuated by the RING domains of RNF43/ZNRF3, which ubiquitylated Fz receptors for degradation. **(b)** The ubiquitylation function of RNF43/ZNRF3 can be quenched by complexing with R-spondin and Lgr4/5/6, which results in more Fz receptors on the cell surface to potentiate Wnt signalling. **(c)** Proposed cartoon model of Norrin mediated signalling with Lgr4/5/6 for neurogenesis and neuroprotection. Unidentified proteins may involve in the signalling. **(d)** Proposed cartoon model of Norrin forming the signalling complexes with Lgr4/5/6 and Fz4–LRP5/6–Tspan12.

Secondly, our complex structure revealed that Norrin dimer binds separately to two copies of Lgr4_ECD_ in a 2:2 stoichiometry (Fig. 2 and Supplementary Fig. S1c, which is similar to the 2:2 complex of Norrin–Fz4_CRD_ structures^33,38^. In contrast, Rspo forms a 1:1 complex with Lgr4/5/6_ECD_ structures^14–16,18,34^. Interestingly, Zebisch and Jones reported that the ternary complex of Rspo2 with Lgr5_ECD_ and ZNRF3_ECD_ revealed a 2:2:2 stoichiometry via the dimerization of ZNRF3_ECD_^37^. Therefore, future studies will need to determine the role of oligomerization of Lgr4/5/6 with divergent ligand recognitions.

Thirdly, our structural analysis revealed that several residues of Norrin for Lgr4 binding are inherited mutation sites (Fig. 6b and Supplementary Table 4). Does Norrin–Lgr4/5/6 signalling axis lead to the pathological progress of ND and FEVR?

Fourthly, our structural comparison of Norrin–Lgr4_ECD_ with Rspo1–Lgr4_ECD_ and Norrin–Fz4_CRD_ revealed several features. The interface of Lgr4 for Norrin binding largely overlaps with the interface for Rspo1 binding (Fig. 5a and 6a). The observation implies a direct competition between Norrin and Rspo1 for Lgr4/5/6 binding. Therefore, it will be interesting to test whether Norrin can serve as an antagonist for Rspo–Lgr4/5/6–Rnf43/Znrf3 signalling axis? As shown in Fig. 5b and 6b, each chain of Norrin only either binds to Fz4 or Lgr4. In other words, Fz4 and Lgr4 compete with each other for Norrin binding. As Norrin is a dimer, does Norrin use one chain for Lgr4 binding and the other chain for Fz4 binding (Fig. 7d)? If so, what is the biological function induced by Norrin–Lgr4–Fz4 signalling axis?

Fifthly, a new paradigm of ligand–receptor recognition of cystine-knot growth factor with Lgr receptor. Lgr1 (follicle-stimulating hormone receptor, FSHR) recognizes the heterodimeric FSH mainly via its concave surface of LRR^46,47^ (Supplementary Fig. S6a). This binding mechanism also occurred in Lgr2 (luteinizing choriogonadotropin hormone receptor, LHCGR) and CG interactions^48^ (Supplementary Fig. S6b). Interestingly, the fashion of the concave surface of LRR for ligand recognition is structural conserved among Lgr1/2 and Lgr4/5/6 for binding to Rspo and Norrin (Fig. 2), although Lgr4/5/6 contains more than six repeats of LRR than Lgr1/2. Moreover, distinct from Norrin using the regions (β1 to β2 and β5 to β6) far from the centre region of dimer assembly for Lgr4 binding and forming a 2:2 complex with Lgr4_ECD_ (Fig. 2), FSH and CG use the centre regions for receptor binding and forming a 2:1 complex with the ECDs of FSHR and FSHR, respectively (Supplementary Fig. S6). Interestingly, previous complex structures of FSH–FSHR_ECD_ have been reported in a dimeric or trimeric state^46,47^; however, a recently determined structure revealed that a CG dimer complex with a monomeric full-length LHCGR and G proteins. Therefore, it will be interesting to dissect whether Lgr receptors would undergo the oligomerization in living cells upon ligand bound and how the distinct states of oligomerization lead to ligand recognition and signalling activation.

Lastly, as most secreted ligands bind to one receptor or several evolutionary related receptors, Norrin serves as a unique paradigm for the interaction with several phylogenetically unrelated receptors of Fz4 and Lgr4 as well as Tetraspanin-12^32,49^. At present, the more biological functions of Norrin mediated signalling pathways through different receptors remain to be determined.

## Methods

### Construct design and cloning

For the Lgr4_ECD_-mVenus construct (Supplementary Fig. S1a), the coding segments of human Lgr4 (UniProt code: Q9BXB1; residues 28-396) with hagfish variable lymphocyte receptor (VLR; UniProt code: Q4G1L2; residues 131-200) C-terminally was cloned into the pHLsec-mVenus-12H vector^33,50^. The SUMO-Norrin construct (Supplementary Fig. S1a) was described previously^33^. For protein-protein interaction experiments, human Lgr5 (UniProt code: O75473) coding segment PCR-amplified from hLgr5_ECD_ pHLsec^37^ was cloned into the pHLsec-Fc-Avi-6H vector^32^, which contains the human immunoglobulin heavy constant gamma 1 followed by an Avi tag that can be biotinylated by BirA ligases and a 6xHis tag. The coding segment of human Lgr6 (Uniprot code: Q9HBX8; residues 31-405) with VLR (UniProt code: Q4G1L2; residues 131-200) C-terminally was cloned into the pHLsec-Fc-Avi-6H vector^32^. The AP-Norrin construct was described previously^29^. Mutants were generated using two-step overlapping PCR experiments. All constructs were confirmed by DNA sequencing.

### Protein expression and purification

HEK293T (ATCC CRL-11268) cells were maintained in a humidified 37° C incubator with 5% CO_2_ in Dulbecco’s Modified Eagle Medium (DMEM, MilliporeSigma) supplemented with 2 mM L-Glutamine (L-Glu, Gibco), 0.1 mM non-essential amino acids (NEAA, Gibco) and 10% [v/v] Fetal Bovine Serum (FBS, Gibco). The FBS concentration was lowered to 2% [v/v] after transfection with the DNA using polyethylenimine (MilliporeSigma 408727) as described previously^51^. Constructs of SUMO-Norrin and Lgr4_ECD_-mVenus were co-transfected in HEK293T cells in the presence of 5 μM of the class I α-mannosidase inhibitor, kifunensine^52^ and 4 mM valproic acid^33^. Conditioned media were collected five days post-transfection and supplemented with 10 mM HEPES, pH 7.5, and 5 mM imidazole. The His-tagged sample was purified by immobilized metal affinity chromatography (IMAC) using Ni Sepharose Excel resin (GE Healthcare Life Sciences). The IMAC eluted sample was dialyzed and treated with His-tagged Human Rhinovirus 3C protease. The treated sample was further purified by IMAC and was subjected to size-exclusion chromatography (SEC) using HiLoad Superdex 200 pg (GE Healthcare Life Sciences) in 10 mM HEPES, pH 7.5, 0.15 M NaCl. For the AP fusion proteins (AP-Norrin), HEK293T cells were grown in 6-well plates and transfected with the DNA using polyethylenimine (MilliporeSigma 408727) as described previously^32,50^. Conditioned media were collected two days post-transfection. For the biotinylated bait preparations, the plasmid DNA mix of target vector and pHLsec-BirA-ER^33^ into a 3 to 1 ratio transfected in HEK293T cells in the presence of 0.1 mM Biotin (MilliporeSigma B4639). The biotinylated baits were purified from conditioned media collected two days post-transfection by the IMAC method.

### Crystallization and data collection

Purified Norrin–Lgr4_ECD_ complex protein was concentrated to 10 mg/ml and was subjected to sitting drop vapour diffusion crystallization trials in 96-well Greiner plates consisting of 100 nl protein solution and 100 nl reservoir using a Cartesian Technologies dispensing instrument^53^. Crystallization plates were stored and imaged via the ROCK IMAGER 1000 at 294 K. Norrin– Lgr4_ECD_ complex crystallized in 0.1 M Bis-Tris propane, pH 7.5, 0.2 M sodium malonate, 20% polyethylene glycol (PEG) 3350. For cryoprotection, crystals were transferred into a reservoir solution supplemented with 25% glycerol and subsequently cryo-cooled in liquid nitrogen. X-ray diffraction data were collected at 100 °K Diamond Light Source (Oxfordshire, UK) at beamlines I24. Diffraction data were indexed, integrated, and scaled using the XIA2 system^54^, coupled with DIALS^55,56^ and POINTLESS^57^. Diffraction anisotropy was further corrected using STARANISO^58^.

### Structure determination and refinement

The Norrin–Lgr4_ECD_ complex structure was determined by molecular replacement in PHASER^59^ using the Norrin structure (PDB ID: 5BQ8) and Lgr4_ECD_ (PDB ID: 4QXE) as the search models. The resulting map showed three copies of Norrin and Lgr4_ECD_ fit into the electron density. The models were completed by manual building in COOT^60^ and refinement was performed using PHENIX Refine^61^ with translation-libration-screw (TLS) parameterization and AutoBUSTER^62^ with the rigid body refinement, TLS parameterization, and geometric restraints of non-crystallographic symmetry and high-resolution reference structures. MOLPROBITY^63^ was used to validate the models. The crystallographic statistics are listed in Table S1.

### Structure analysis

Structure-based multiple sequence alignment was generated using ClustalΩ^64^ and ESPript^65^. Structure superposition was performed using SUPERPOSE^66^ in the CCP4 suite^67^. Electrostatic potential calculations were generated using APBS tools^68^ and interface areas of proteins were analyzed with the PISA webserver^69^. High-quality images of the molecular structures were generated with the PyMOL Molecular Graphic System (Version 2.5, Schrödinger, LLC). Schematic figures and other illustrations were prepared using GraphPad Prism (GraphPad Software, LLC) and Corel Draw (Corel Corporation).

### Alkaline Phosphatase based binding assay

The biotinylated baits were immobilized on 96-well streptavidin-coated plates (Thermo Fisher Scientific 15500). The wells were then washed three times with wash buffer (10 mM HEPES, pH 7.5, 0.15 M NaCl, 0.05% (w/v) Tween-20) and incubated with a 10-fold dilution of Blocker bovine serum albumin (BSA; Thermo Fisher Scientific 37525) in wash buffer for 1 h at 25 °C.

The wells were washed with wash buffer and incubated with conditioned media containing AP-Norrin probes proteins for 2 h at 25 °C. The wells were subsequently washed three times with wash buffer and incubated with BluePhos phosphatase substrate solution (Kirkegaard and Perry Laboratories 50-88-00) to visualize the bound AP probes.

## Acknowledgements

We thank Diamond Light Source for beamtime and the staff at beamline I24 for assistance with data collection. Margaret Jones and Tom Walter for technical support. Weixian Lu and Yuguang Zhao for help with tissue culture. XXX for comments on the manuscript. Jeremy Nathans (Johns Hopkins University) for the gift of AP-Norrin construct. Clemens Vonrhein (Global Phasing Ltd) for the advice on data processing. This work was supported by grants to EYJ from XXX. The WTCHG is supported by the Wellcome Trust (XXX). TC was supported by the Human Science Frontier Program Organization Fellowship (LT000130/2017-L).

## Author contributions

Conceptualization: TC, EYJ. Methodology: TC. Investigation TC, KH, FH. Data analysis and visualization: TC, FH, EYJ. Funding acquisition: TC, EYJ. Supervision: TC, EYJ. Writing: TC, EYJ.

## Additional information

Coordinates and structure factors have been deposited in the Protein Data Bank (PDB) with an accession code XXXX.

