## Supplemental data for "Structural basis of Norrin recognition by Lgr4/5/6"

Running Title: Structure-function of Norrin and Lgr4/5/6

Keywords: Wnt signalling, angiogenesis, blood-brain barrier, neuroprotection, retinal vascular disease, Norrin, Norrie disease

### Supplemental data

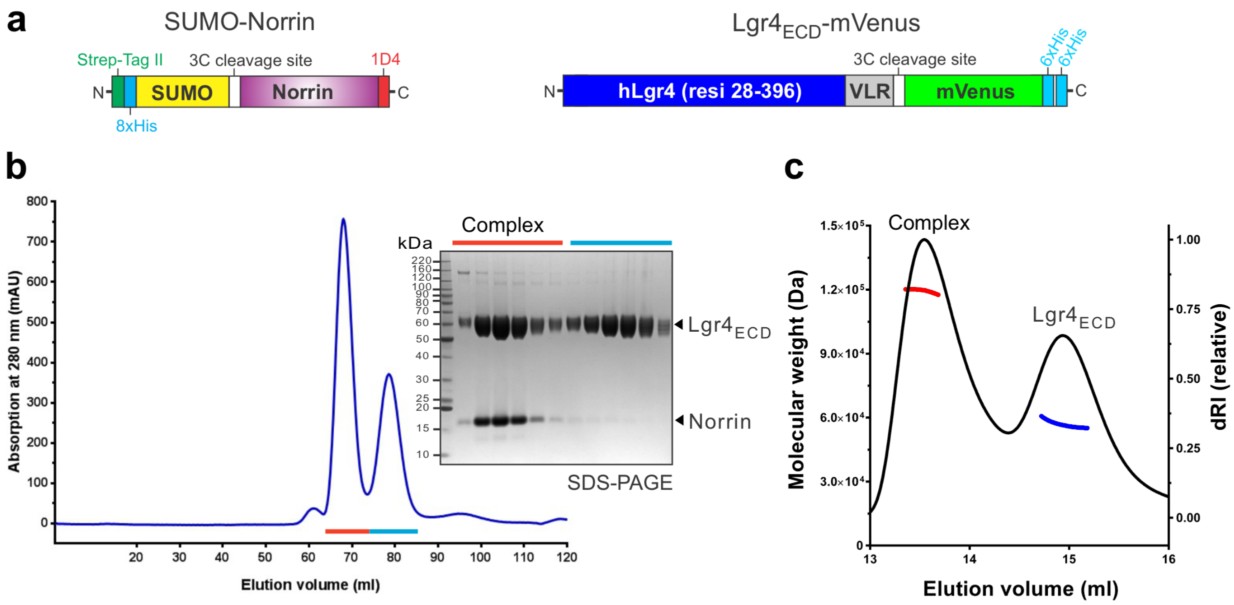

**Supplementary Figure 1 | Protein purification and biophysical analysis of the Norrin–Lgr4$_{ECD}$ complex.**
**(a)** Schematic diagrams of the expression constructs of SUMO-Norrin and Lgr4$_{ECD}$-mVenus. See also Materials and Methods for detailed information. **(b)** SEC elution profile and SDS-PAGE of Norrin–Lgr4$_{ECD}$ complex and Lgr4$_{ECD}$ under reducing conditions with fractions analysed marked by red and cyan lines, respectively. **(c)** Analysis of the formation of a 2:2 complex of dimeric Norrin bound to two copies of monomeric Lgr4$_{ECD}$ by SEC-MALS. The profiles of molecular weight (left ordinate axis) of Norrin–Lgr4$_{ECD}$ complex and monomeric Lgr4$_{ECD}$ are shown as red and blue lines, respectively. The profile of the differential refractive index (right ordinate axis) is shown as a black line.

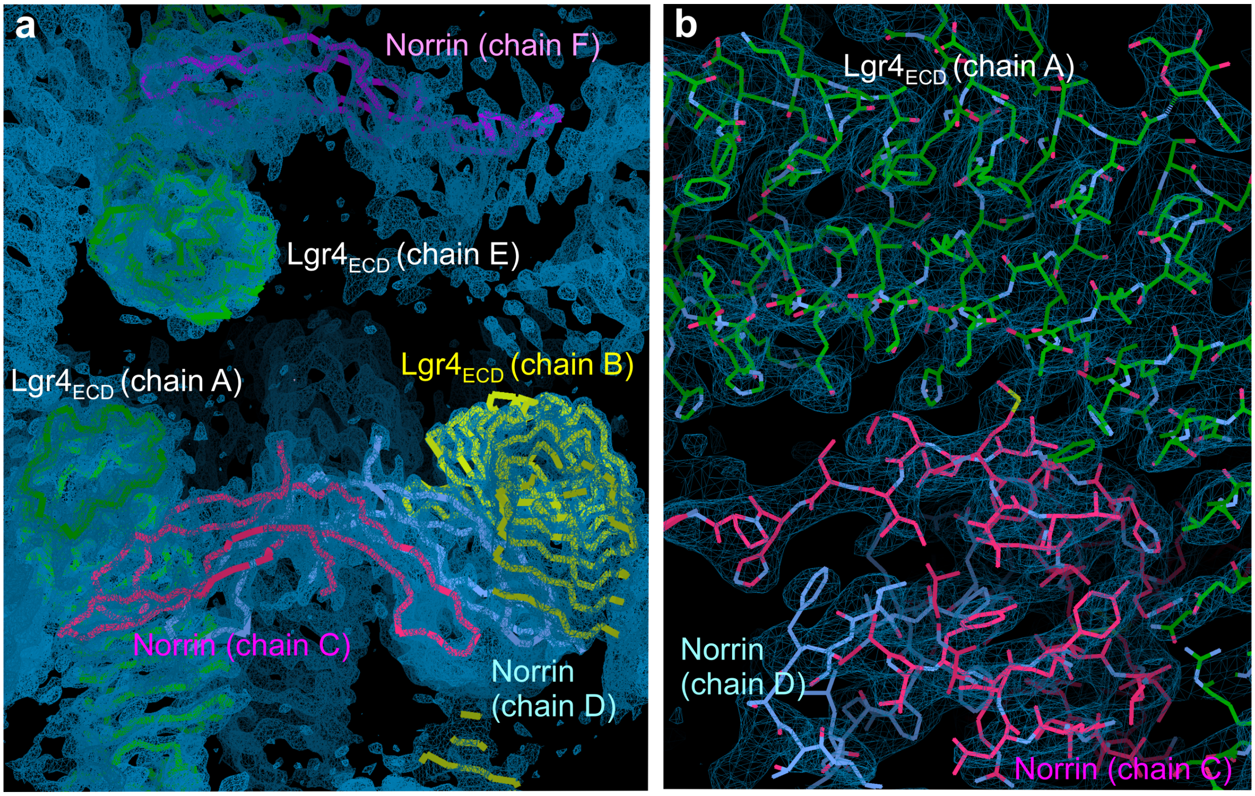

**Supplementary Figure 2 | Electron density of Norrin–Lgr4$_{ECD}$.**
**(a)** The overall sigmaA-weighted 2|$F_O$|-|$F_C$| electron density map after refinement is contoured at the 1.3 σ level as blue meshes for all chains of Norrin and Lgr4$_{ECD}$ from the asymmetric unit. **(b)** A view of the electron density for the binding interface in the Norrin–Lgr4$_{ECD}$ complex is shown as the sigmaA-weighted 2|$F_O$|-|$F_C$| map (blue meshes), contoured at the 1.0 σ level.

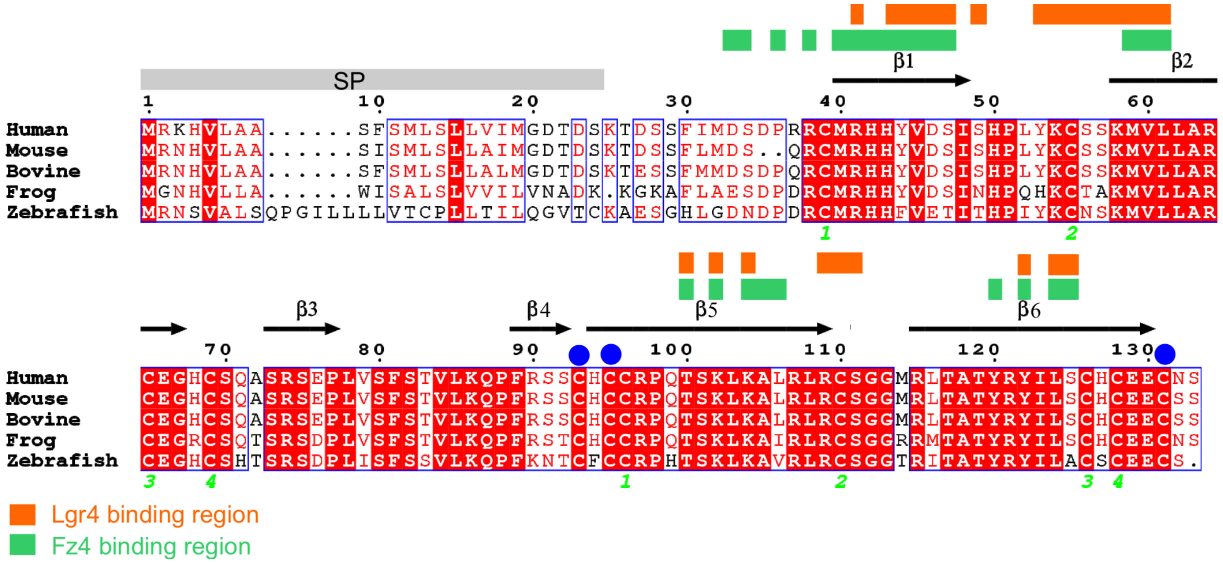

**Supplementary Figure 3 | Sequence alignment of Norrin proteins.**
Protein sequences of Norrin from human (UniPort code: Q00604), mouse (UniPort code: P48744), bovine (UniPort code: Q2KI78), frog (UniPort code: C0IW62), and zebrafish (UniPort code: E7F073) are aligned by ClustalΩ. Above the alignment is the secondary structure assignment of Norrin. Orange and green rectangles highlight residues for Lgr4 binding and Fz4 binding, respectively. The region of signal peptide (SP) is marked with a grey rectangle. Intramolecular disulphide bonds are numbered and three intermolecular disulphide bonds are marked with blue circles.

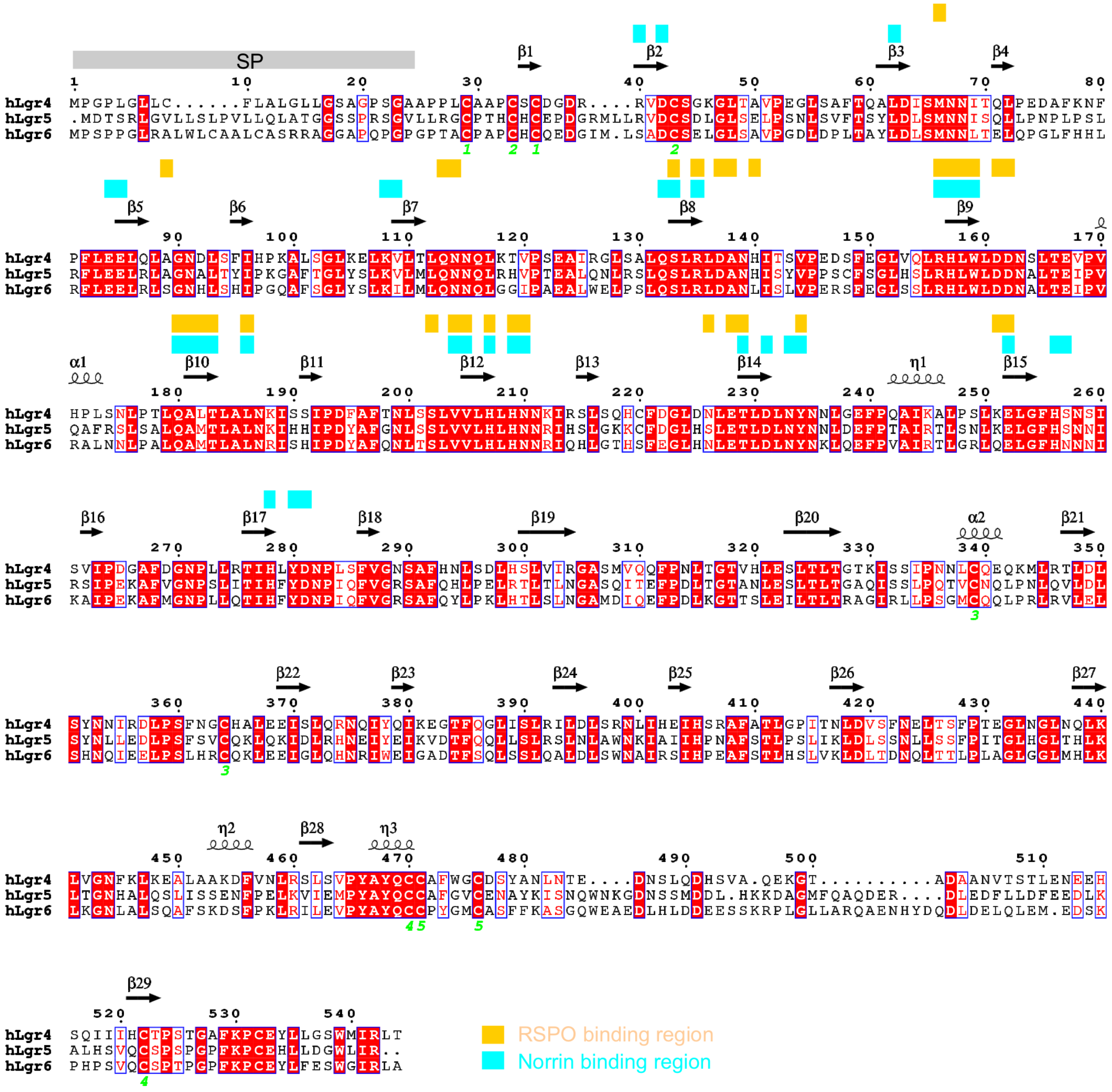

**Supplementary Figure 4 | Analysis of Lgr4 and Lgr5/6 homologues.**
An alignment of the sequence of human Lgr4 (UniPort code: Q9BXB1) with Lgr5 (UniPort code: O75473) and Lgr6 (UniPort code: Q9HBX8). Above the alignment is the secondary structure assignment of Lgr4, derived from the structure (PDB code 4KT1). Gold and cyan rectangles highlight residues for RSPO binding and Norrin binding, respectively. The region of signal peptide (SP) is marked with a grey rectangle. Disulphide bonds of Lgr4 are numbered.

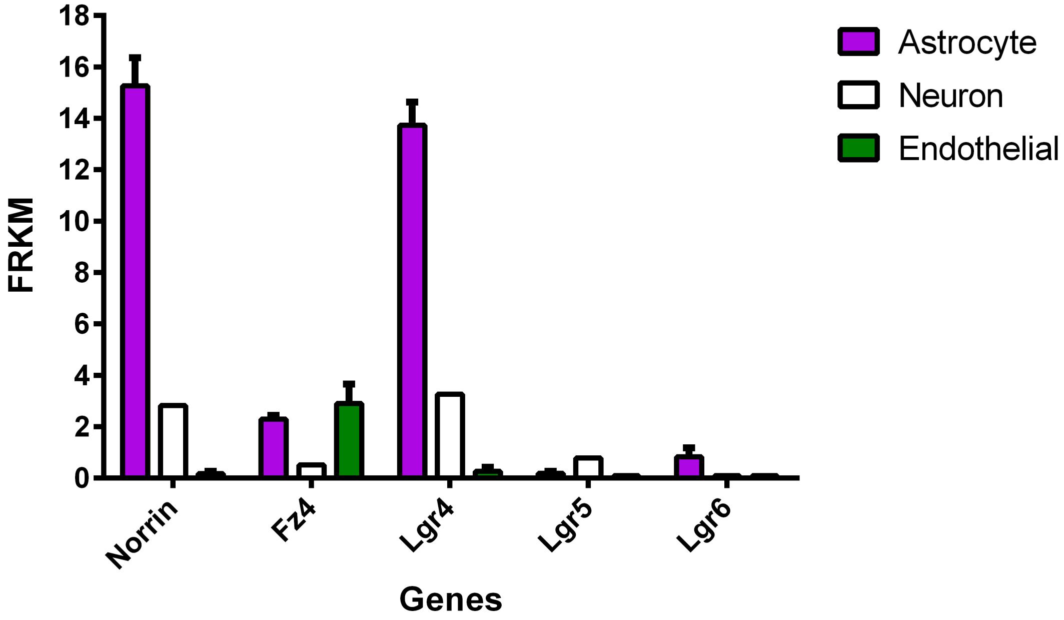

**Supplementary Figure 5 | Human brain RNA-seq data of gene transcripts of Norrin, Fz4, and Lgr4/5/6.**
Expression levels (Fragments Per Kilobase Million, FPKM) are based on RNA-seq of purified cells from human brains by the immunopanning method. RNA-seq expression plots were rendered with the database of Brain RNA-Seq (https://www.brainrnaseq.org).

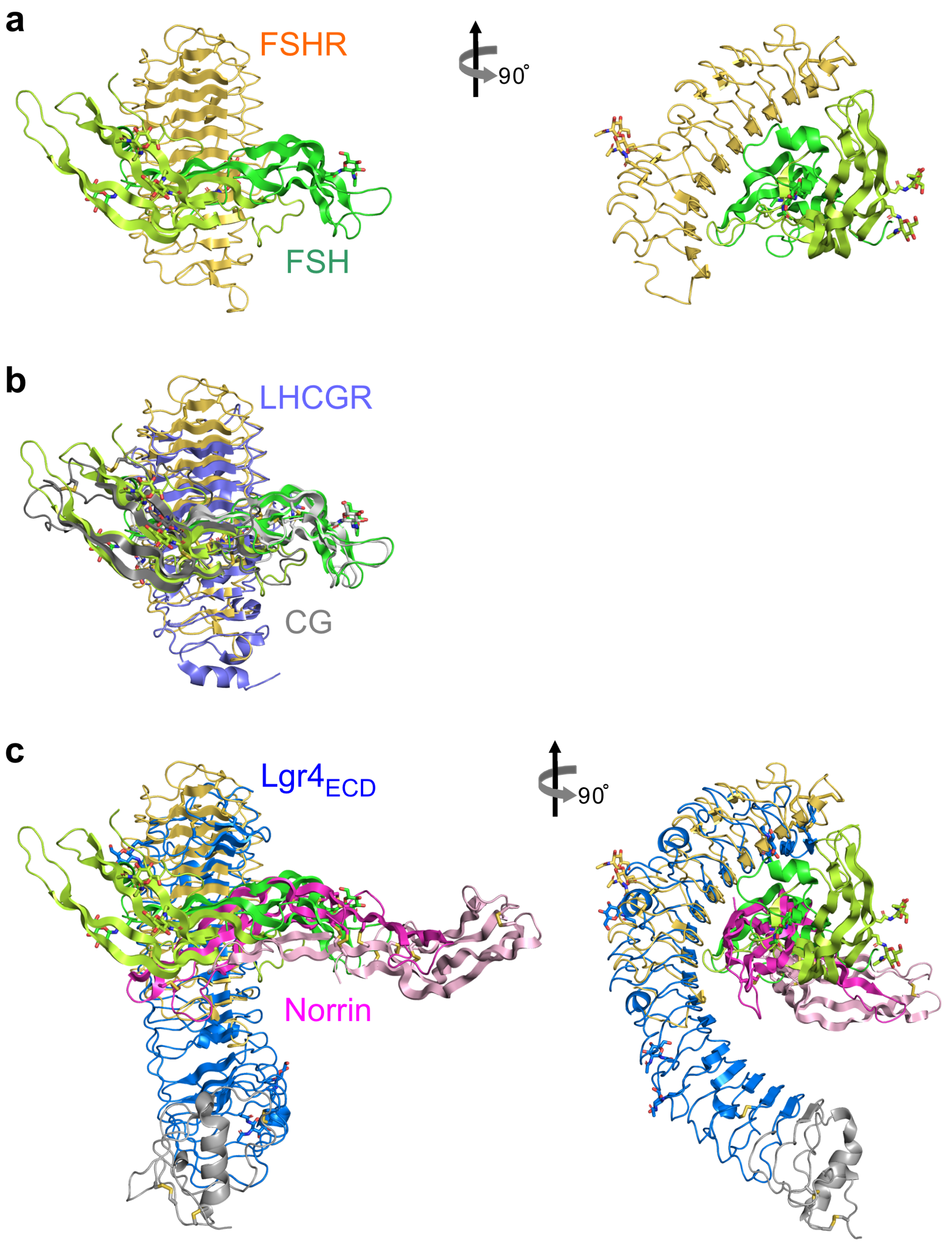

**Supplementary Figure 6 | Structural comparison of the ligand and receptor complexes.** **(a)** The complex structure of FSH–FSHR (PDB code: 1XWD). FSH is composed of α-chain (green) and β-chain (lemon) and FSHR is coloured gold. N-linked glycans are shown as green sticks. **(b)** Structural comparison of FSH–FSHR with CG–LHCGR (PDB code: 7FIH). CG is composed of α-chain (light grey) and β-chain (dark grey) and LHCGR is coloured slate. **(c)** Superpositions of Norrin (pink and magenta)–Lgr4 (blue) with FSH–FSHR and CG–LHCGR based on receptor parts.

**Supplementary Table S1. Molecular properties of the proteins determined by SEC-MALS.**

| Protein[a] | Number of N-glycan sites | $MW_{Theoretical}$ (kDa)[b] | $MW_{Measured}$ (KDa) |
|---|---|---|---|
| $Lgr4_{ECD}$ | 3 | 55.6 (monomer) | 56.4 ± 0.4 |
| Norrin– $Lgr4_{ECD}$ | 6 (2:2 complex) | 138.3 (2:2 complex) | 119.8 ± 0.7 |

[a]The proteins were produced from HEK293T cells in the presence of the N-glycosylation processing inhibitors, kifunensine resulting in limited glycosylation and were treated with endoglycosidase-$F_1$.
[b]The measured molecular weight ($MW_{Measured}$) is in general agreement with theoretical molecular weight ($MW_{Theoretical}$) predicated based on the primary sequence plus the molecular weight of N-linked glycans (see Materials and Methods of SEC-MALS analysis for detailed information of calculation).

**Supplementary Table S2. Data collection and refinement statistics.**

| | Norrin–$Lgr4_{ECD}$ [a] |
|---|---|
| **X-ray Source** | DLS I24 |
| **Data collection** | |
| Space group | *C*2 |
| Unit-cell parameters | |
| *a*, *b*, *c* (Å) | 121.34, 113.96, 199.46 |
| $\alpha$, $\beta$, $\gamma$ (°) | 90.00, 97.51, 90.00 |
| No. of crystals | 1 |
| Wavelength | 1.07198 |
| Resolution (Å) [b] | 66.46 - 3.05 (3.45 - 3.05) |
| No. of unique reflections [b] | 29160 (1458) |
| Completeness (%) [b] | 93.0 (68.1) |
| Multiplicity[b] | 6.6 (5.7) |
| I / σ (*I*) [b] | 6.7 (1.6) |
| $R_{pim}$ (%) [b,c] | 11.0 (77.9) |
| $CC_{1/2}$ (%) [b,d] | 99.6 (44.0) |
| **Refinement** | |
| Resolution (Å) | 49.44 - 3.05 |
| $R_{work}$ / $R_{free}$ | 25.04 / 29.04 |
| No. atoms | |
| Protein | 12514 |
| Ligand | 154 |
| *B*-factors | |
| Protein | 124.04 |
| Ligand | 142.86 |
| R.m.s deviations [e] | |
| Bond lengths (Å) | 0.01 |
| Bond angles (°) | 1.54 |
| Ramachandran plot | |
| Favored (%) | 95.24 |
| Allowed (%) | 4.76 |
| PDB code | |

[a] Diffraction anisotropy correction yield an ellipsoidal resolution boundary with limits of 3.05, 4.07, 3.88 Å along the a*, b*, and c* axes, respectively.
[b] Values in parentheses are for highest-resolution shell.
[c] $R_{pim}$: Precision-indicating merging R-factor.
[d] $CC_{1/2}$: correlation coefficients between random half data sets.
[e] R.m.s deviations: root mean square deviation from ideal geometry.

**Supplementary Table S3. The electrostatic interactions between Norrin with Lgr4$_{ECD}$.** Resides of Norrin associated with disease mutations are highlighted in red.

| Norrin Residue | Group | Lgr4$_{ECD}$ Residue | Group | Interaction |
|---|---|---|---|---|
| Agr41 | side chain NH2 | Glu85 | side chain OE1 | Salt bridge |
| His43 | side chain NE2 | Glu85 | side chain OE1 | Salt bridge |
| His43 | main chain O | Agr156 | side chain NH1 | Hydrogen bond |
| His43 | main chain O | Agr156 | side chain NH2 | Hydrogen bond |
| Lys54 | side chain NZ | Asp281 | side chain OD1 | Salt bridge |
| Lys54 | side chain NZ | Asp281 | side chain OD2 | Salt bridge |
| Cys55 | main chain N | Typ234 | side chain OH | Hydrogen bond |
| Ser56 | side chain OG | Asn210 | side chain ND2 | Hydrogen bond |
| Ser57 | side chain OG | His207 | side chain NE2 | Hydrogen bond |
| Ser57 | side chain OG | His209 | side chain NE2 | Hydrogen bond |
| Met59 | main chain O | Trp159 | side chain NE1 | Hydrogen bond |
| Lys102 | side chain NZ | Asp63 | side chain OD2 | Hydrogen bond |
| Agr109 | side chain NH1 | Asp162 | side chain OD2 | Hydrogen bond |
| Leu124 | main chain O | Arg40 | side chain NH1 | Hydrogen bond |

**Supplementary Table S4**. **Classification of Norrin functional domains and disease-associated mutations.**

The Norrin sequence (residues 25-133; mature form by excluding the signal peptide) is used for analyses and categorization (including dimer interface, Fz4 binding site, HSPGs binding site and Lgr4 binding site). Norrin disease-association mutations (including missense, frameshift, nonsense) are identified from retinal disorders including Norrie Disease, Familial Exudative Vitreoretinopathy, Retinopathy Of Prematurity, Coats' disease, Persistent Fetal Vasculature, Primary Retinal Dysplasia, Venous Insufficiency, and Primary Hyperplastic Persistent Vitreous. The information on Norrin disease-association mutations were summarized from Nikopoulos et al., 2010b and two homepages ( http://www.uniprot.org/uniprot/Q00604 and http://www.medmolgen.uzh.ch/research/eyediseases/norriedisease/Norrinmutations.html). Notably, cystine residues are colored in red.

| Residues | Dimer interface | Fz4 binding site | HSPGs binding site | Lgr4 Binding site | Disease-association mutations |
|---|---|---|---|---|---|
| LYS 25 | | | | | |
| THR 26 | | | | | |
| ASP 27 | | | | | |
| SER 28 | | | | | |
| SER 29 | | | | | S29X |
| PHE 30 | | | | | |
| ILE 31 | | | | | |
| MET 32 | | | | | |
| ASP 33 | | Fz4 binding | | | |
| SER 34 | | Fz4 binding | | | |
| ASP 35 | | | | | D35fs |
| PRO 36 | | Fz4 binding | | | |
| ARG 37 | | | | | |
| ARG 38 | | Fz4 binding | | | R38C |
| CYS 39 | | | | | C39R |
| MET 40 | | Fz4 binding | | | |
| ARG 41 | | Fz4 binding | | Lgr4 binding | R41K R41S R41T |
| HIS 42 | | Fz4 binding | | | H42E H42R |
| HIS 43 | | Fz4 binding | | Lgr4 binding | H43Q H43R H43fs |
| TYR 44 | Dimer | Fz4 binding | | Lgr4 binding | Y44C Y44X |
| VAL 45 | | Fz4 binding | | Lgr4 binding | V45E V45M V45fs |
| ASP 46 | | Fz4 binding | | Lgr4 binding | |
| SER 47 | | Fz4 binding | | Lgr4 binding | |
| ILE 48 | Dimer | | | | I48fs |
| SER 49 | Dimer | | | Lgr4 binding | |
| HIS 50 | Dimer | | | | H50D |
| PRO 51 | Dimer | | | | |
| LEU 52 | | | | | |
| TYR 53 | | | | Lgr4 binding | |
| LYS 54 | | | | Lgr4 binding | K54N |
| CYS 55 | | | | Lgr4 binding | C55R C55F |
| SER 56 | | | | Lgr4 binding | |
| SER 57 | | | | Lgr4 binding | S57X |
| LYS 58 | | | HSPGs binding | Lgr4 binding | K58N |
| MET 59 | | Fz4 binding | | Lgr4 binding | |
| VAL 60 | | Fz4 binding | | Lgr4 binding | V60E |
| LEU 61 | | Fz4 binding | | Lgr4 binding | L61F L61I L61P |
| LEU 62 | Dimer | | | | |
| ALA 63 | Dimer | | | | A63D A63S |
| ARG 64 | Dimer | | | | |
| CYS 65 | Dimer | | | | C65W C65Y |
| GLU 66 | Dimer | | | | |
| GLY 67 | Dimer | | | | G67E G67R |

| | | | | | | | | |
|---|---|---|---|---|---|---|---|---|
| HIS 68 | Dimer | | | | | | | |
| CYS 69 | Dimer | | | | C69S | C69fs | | |
| SER 70 | Dimer | | | | | | | |
| GLN 71 | Dimer | | | | | | | |
| ALA 72 | Dimer | | | | | | | |
| SER 73 | Dimer | | | | S73X | | | |
| ARG 74 | Dimer | | | | R74C | | | |
| SER 75 | Dimer | | | | S75C | S75P | | |
| GLU 76 | Dimer | | | | | | | |
| PRO 77 | Dimer | | | | | | | |
| LEU 78 | Dimer | | | | | | | |
| VAL 79 | Dimer | | | | | | | |
| SER 80 | Dimer | | | | | | | |
| PHE 81 | Dimer | | | | | | | |
| SER 82 | | | | | | | | |
| THR 83 | Dimer | | | | | | | |
| VAL 84 | Dimer | | | | | | | |
| LEU 85 | Dimer | | | | | | | |
| LYS 86 | | | | | | | | |
| GLN 87 | | | | | | | | |
| PRO 88 | Dimer | | | | | | | |
| PHE 89 | Dimer | | | | F89L | F89fs | | |
| ARG 90 | | | | | R90C | R90P | | |
| SER 91 | Dimer | | | | | | | |
| SER 92 | | | | | S92P | | | |
| CYS 93 | Dimer | | | | C93R | | | |
| HIS 94 | Dimer | | | | H94fs | | | |
| CYS 95 | Dimer | | | | C95F | C95R | C95Y | C95X |
| CYS 96 | Dimer | | | | C96W | C96Y | C96fs | |
| ARG 97 | Dimer | | | | R97P | | | |
| PRO 98 | Dimer | | | | P98L | | | |
| GLN 99 | | | | | | | | |
| THR 100 | | Fz4 binding | | Lgr4 binding | | | | |
| SER 101 | Dimer | | | | S101F | | | |
| LYS 102 | | Fz4 binding | | Lgr4 binding | | | | |
| LEU 103 | Dimer | | | | L103V | | | |
| LYS 104 | | Fz4 binding | | Lgr4 binding | K104N | K104Q | | |
| ALA 105 | | Fz4 binding | | | A105T | | | |
| LEU 106 | | Fz4 binding | | | | | | |
| ARG 107 | | | HSPGs binding | | | | | |
| LEU 108 | Dimer | | | | L108P | | | |
| ARG 109 | | | HSPGs binding | Lgr4 binding | R109X | | | |
| CYS 110 | | | | Lgr4 binding | C110G | C110R | C110S | C110X |
| SER 111 | | | | Lgr4 binding | | | | |
| GLY 112 | | | | | G112E | | | |

| | | | | | | | | |
|---|---|---|---|---|---|---|---|---|
| GLY 113 | | | | | G113D | | | |
| MET 114 | | | | | | | | |
| ARG 115 | | | HSPGs binding | | R115L | | | |
| LEU 116 | Dimer | | | | | | | |
| THR 117 | Dimer | | | | | | | |
| ALA 118 | Dimer | | | | A118D | | | |
| THR 119 | Dimer | | | | | | | |
| TYR 120 | Dimer | Fz4 binding | | | Y120C | Y120X | | |
| ARG 121 | Dimer | | | | R121G | R121L | R121W | R121Q |
| TYR 122 | Dimer | Fz4 binding | | Lgr4 binding | | | | |
| ILE 123 | Dimer | | | | I123N | | | |
| LEU 124 | | Fz4 binding | | Lgr4 binding | L124F | | | |
| SER 125 | | Fz4 binding | | Lgr4 binding | | | | |
| CYS 126 | | | | | C126S | C126R | C126X | |
| HIS 127 | | | | | | | | |
| CYS 128 | | | | | C128R | C128X | | |
| GLU 129 | | | | | | | | |
| GLU 130 | | | | | | | | |
| CYS 131 | Dimer | | | | | | | |
| ASN 132 | Dimer | | | | N132fs | | | |
| SER 133 | | | | | | | | |
